# Evaluation Of Suitability Analysis of Gangetic Water from Upper, Middle, And Lower Ganga Rivers

**DOI:** 10.1101/2024.02.28.582642

**Authors:** Acharya Balkrishna, Sourav Ghosh, Ashwani Nagose, Divya Joshi, Shelly Singh, Kumud, Aditi Saxena, Sanyam Taneja, Vedpriya Arya

**Affiliations:** Patanjali research Foundation, Haridwar-249407, India

**Keywords:** Ganges, Water, Pollution, Industry, Irrigation, Drinking Water, Indo-Gangetic Region

## Abstract

This study assesses the suitability of water from distinct segments of the Ganga River in India for drinking, irrigation, and industrial purposes, utilizing various indices. Conducted in MAR-MAY 2023, water samples were collected from multiple sites along the river, and categorized into upper, middle, and lower stretches. The findings revealed differing levels of suitability for various purposes among the sampled sites within each stretch. Parameters for drinking water quality generally adhered to specified limits across all stretches, except for E. coli, emphasizing the importance of disinfection measures. Moreover, the study evaluated irrigation suitability, with most sites falling within the good to excellent range, albeit some exhibiting elevated sodium levels, particularly in the lower stretch. Analysis of major ion abundance revealed an alkaline nature dominated by sodium and potassium ions over calcium and magnesium ions across all stretches. Heavy metal concentrations were either within specified limits or absent, indicating minimal industrial pollution. However, caution is warranted regarding industrial applications due to water aggressiveness, as suggested by indices such as the Langelier saturation index and Ryznar stability index, which hint at potential corrosion issues. Water quality in the Ganga River in India exhibited variability across the upper, middle, and lower stretches, with each segment presenting unique characteristics and suitability levels. Continuous monitoring and control measures, especially concerning the most probable number (MPN) of contaminants, are imperative to ensure the sustainability of water quality and meet the diverse usage needs along the river. The evaluation of Gangetic water from the Upper, Middle, and Lower Ganga Rivers is crucial for understanding water quality in one of India’s vital river basins. This study assesses physical, chemical, and biological parameters to determine suitability for drinking, irrigation, and aquatic life. Factors like pH, dissolved oxygen, turbidity, nutrients, heavy metals, and microbial load are analyzed to reveal water quality and potential risks. Temporal and spatial variations, influenced by natural processes and human activities, are explored. The findings aid in sustainable water management and conservation in the Gangetic basin, addressing the urgent need to safeguard this critical resource.

## Introduction

As is often known, water stands as the most vital resource necessary for sustaining life on our planet and the Ganga River basin is an important groundwater aquifer recharge zone. Groundwater extraction influences river flow, impacting both its ecological health and downstream water availability. Conversely, variations in river flow can affect groundwater levels and quality, amplifying pollution risks. Contaminants from various sources infiltrate groundwater, threatening human health and ecosystems. One of the primary contributors to water scarcity is inefficient water management practices, leading to over-extraction and depletion of groundwater reserves. Groundwater, a crucial source of water for agriculture, industry, and domestic use, is being exploited at unsustainable rates in many parts of the country. With vast expanses of arable land reliant on irrigation for crop production, groundwater irrigation has become indispensable, especially in regions where surface water sources are limited or unreliable.

Moreover, the Ganga’s flow fuels numerous industrial processes, driving economic growth and development. Thus, the quality of water from the Ganga River holds immense importance, affecting not only the health and well-being of communities but also the productivity of agricultural sectors and the viability of industrial operations (Paul & Sinha 2013). Accounting for 25% of India’s water resources, the Ganga ranks as the thirtieth longest river globally, covering a basin area of 861,404 square kilometers 5 (Rahaman 2009). The Ganga basin represents one of the world’s most densely populated regions, with an average density of 520 persons per square kilometer (Das & Tamminga, 2012). Supporting more than 300 million people across India, Nepal, and Bangladesh, the basin is of immense importance (Gopal, B. 2000). This river system drains around one-fourth of the Indian subcontinent.

Industrialization, urbanization, and agricultural activities have significantly impacted the water quality of the Ganga, raising concerns regarding its suitability for various purposes such as drinking, irrigation, and industrial usage. This paper delves into the comprehensive evaluation of water quality across different stretches of the Ganga, namely the Upper, Middle, and Lower stretches, focusing on 26 water samples collected from states including Uttarakhand, Uttar Pradesh, Bihar, Jharkhand, and West Bengal. The population residing in cities along the Ganga has witnessed exponential growth in recent years, exacerbating the pressure on water resources. For instance, cities like Haridwar, Kanpur, and Patna have experienced substantial increases in population due to factors such as rural-urban migration, industrial development, and natural population growth. Haridwar, nestled in Uttarakhand, is a major pilgrimage site, drawing millions of visitors annually, while Kanpur, located in Uttar Pradesh, is renowned for its industrial prowess, with a significant portion of its population engaged in manufacturing activities. Similarly, Patna, the capital city of Bihar, has witnessed steady population growth. The suitability of Ganga River water for drinking purposes is assessed through the National Sanitation Foundation Water Quality Index (NSF-WQI). These indices serve as vital tools for discerning the safety and purity of water intended for consumption, offering insights into the prevalence of pollutants and their potential health risks. For irrigation purposes, the Sodium Adsorption Ratio (SAR) and Bicarbonate emerge as key indicators of water quality suitability. SAR evaluates the potential sodium hazards in irrigation water, offering crucial insights into soil permeability and fertility. Meanwhile, Kelly’s ratio provides valuable information on the comparative concentrations of essential cations, guiding agricultural practices toward sustainable water usage and soil management. In the realm of industrial water usage, the Langelier Saturation Index (LSI) take precedence in assessing water quality suitability.

## 2. Material Methods

### 2.1. Site Selection

A total of 26 sites have been carefully selected along the banks of the Ganga River. Out of these, five sites are situated in Uttarakhand, while Uttar Pradesh boasts 10 sites. Bihar contributes five sites, Jharkhand has one, and West Bengal is home to five sites along the river. We have segmented the entire expanse of the Ganga River, spanning from Gomukh to Ganga-Sagar, into three distinct sections. According to the altitude, the upper Stretch encompasses sampling sites from Gomukh to Narora, the Middle Stretch encompasses sites from Budaun to Ballia, and the Lower Stretch encompasses sites from Revelganj to Diamond Harbour (Table).

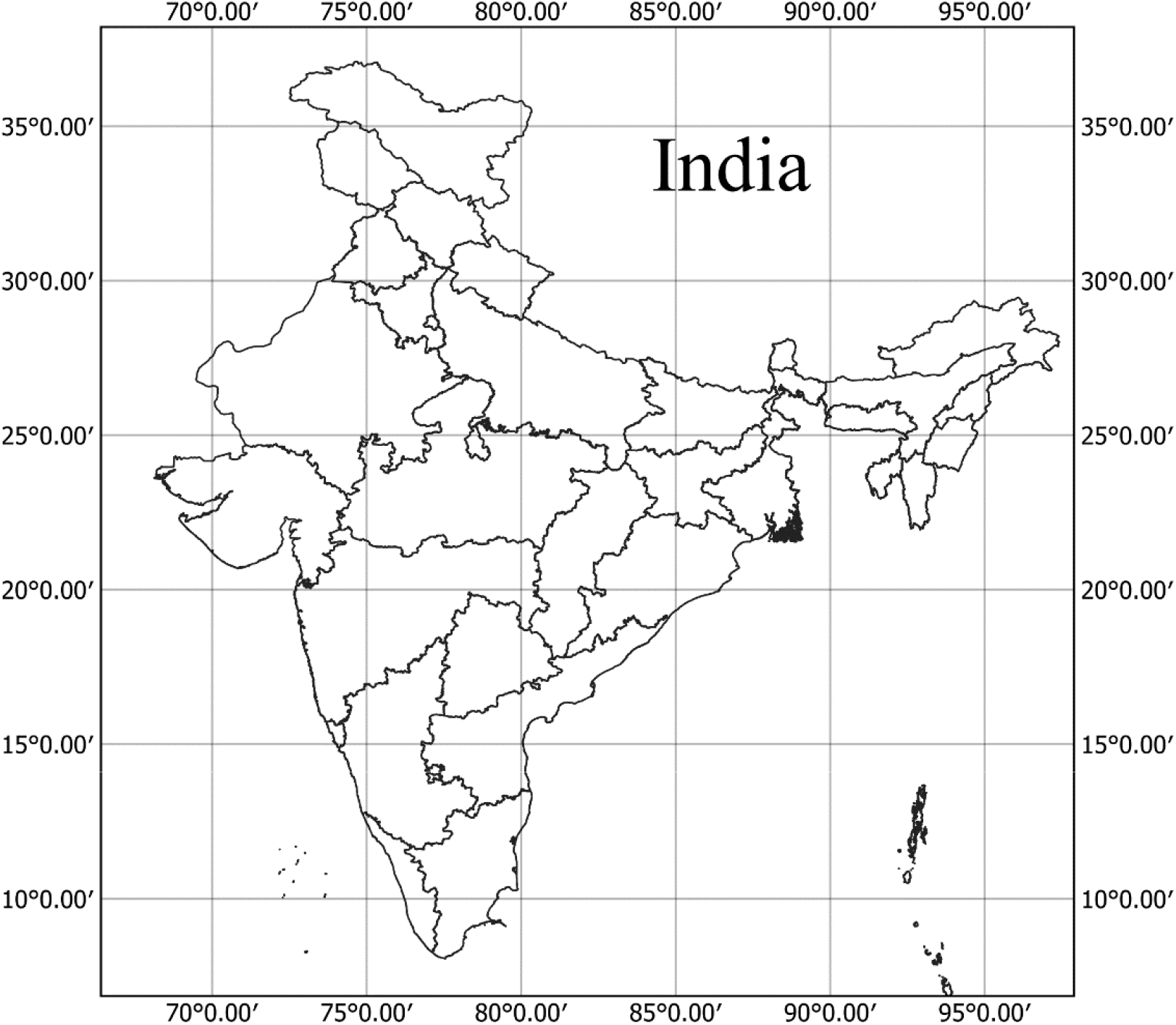

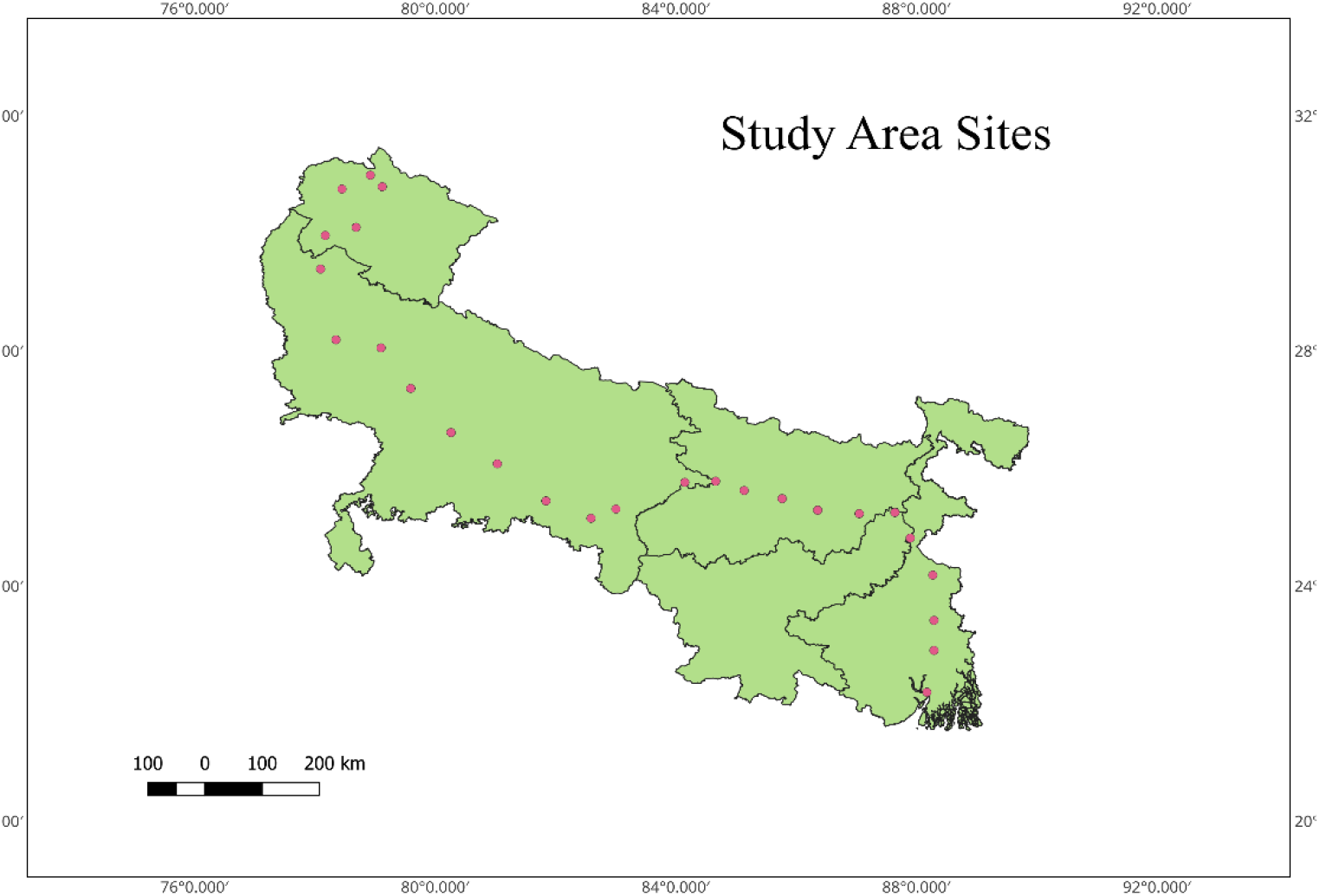

The sites selected for the sampling of water samples have been furnished in Table 1.

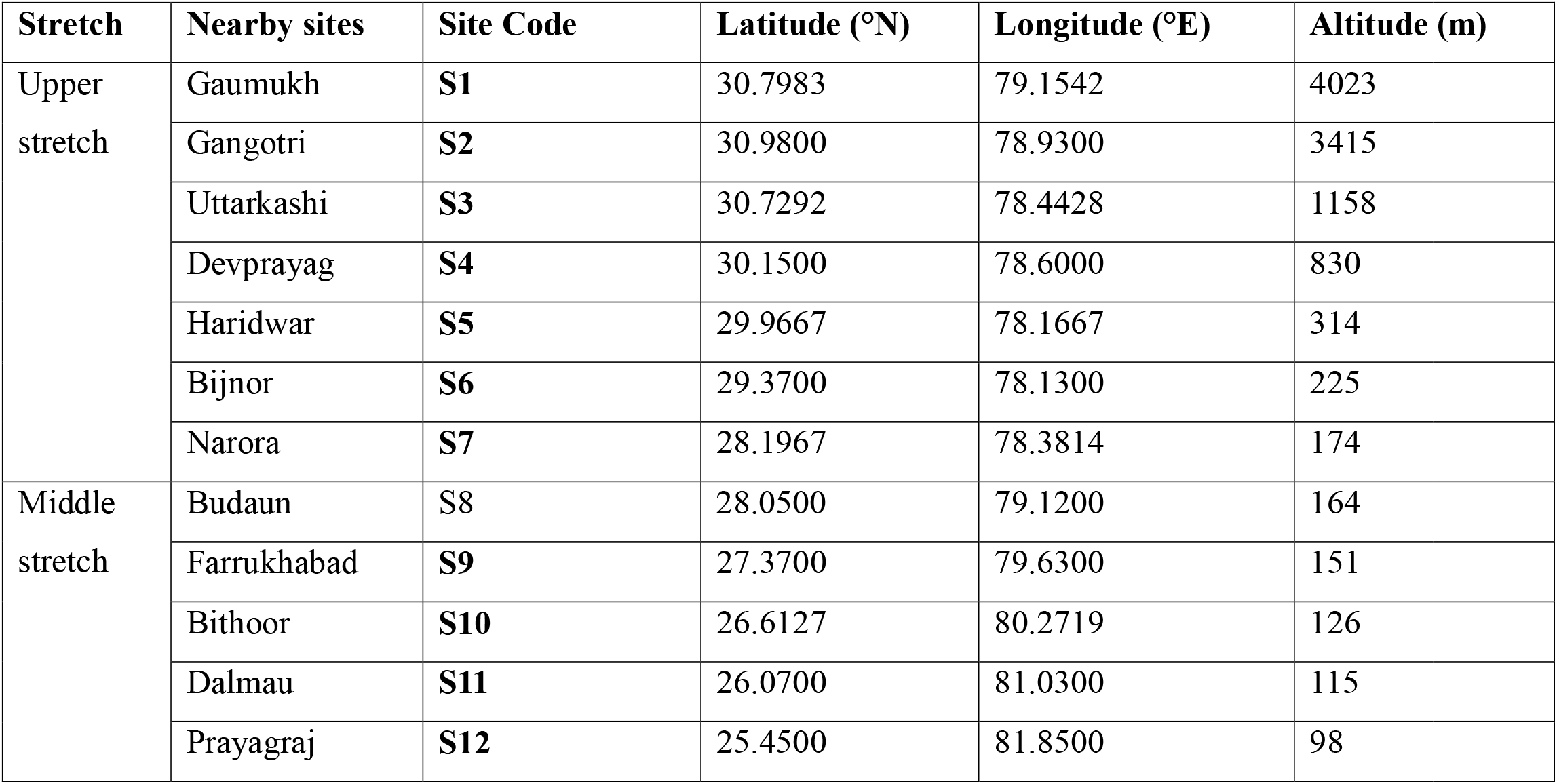
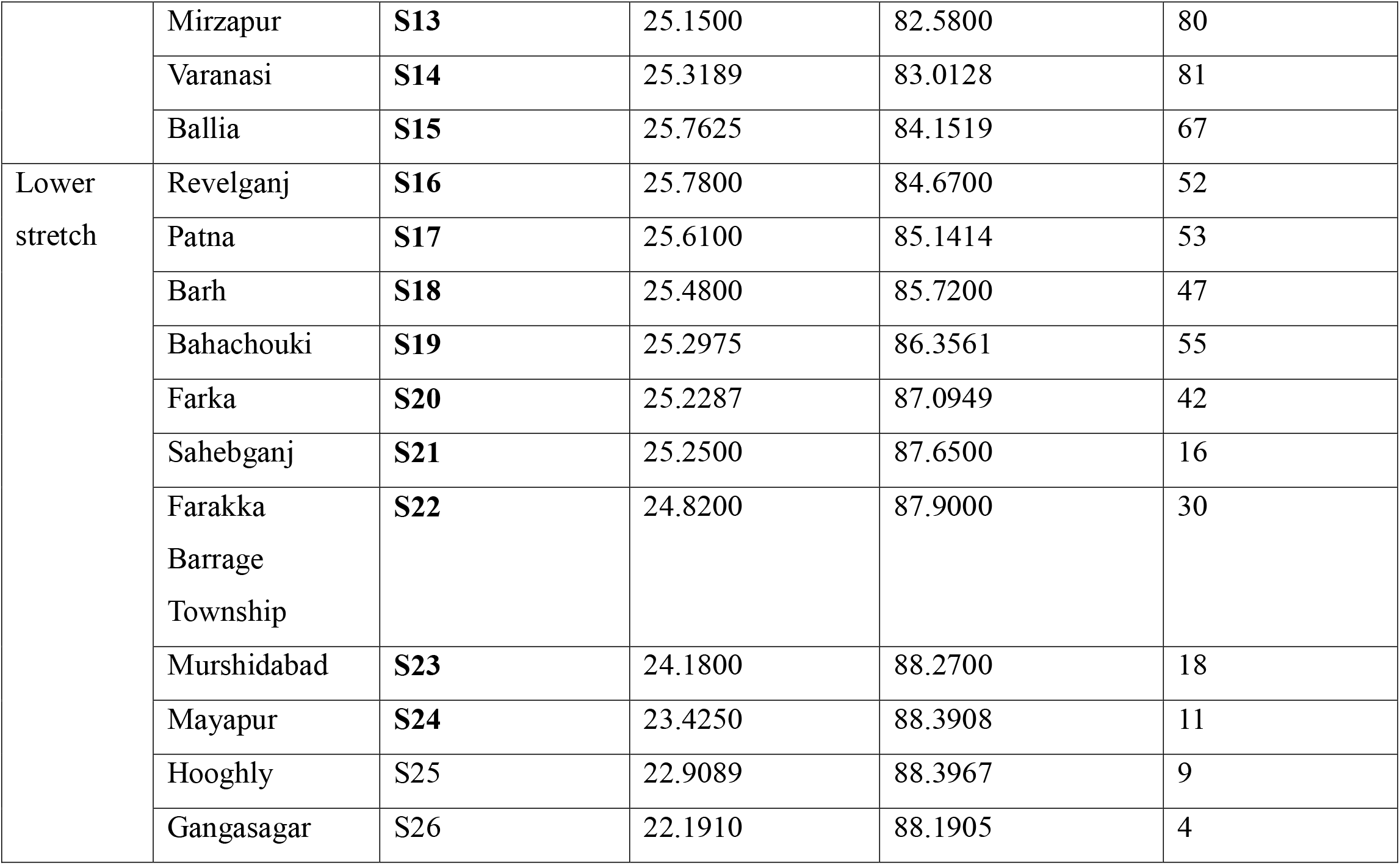

| Stretch | Nearby sites | Site Code | Latitude (°N) | Longitude (°E) | Altitude (m) |
| --- | --- | --- | --- | --- | --- |
| Upper stretch | Gaumukh | **S1** | 30.7983 | 79.1542 | 4023 |
|  | Gangotri | **S2** | 30.9800 | 78.9300 | 3415 |
|  | Uttarkashi | **S3** | 30.7292 | 78.4428 | 1158 |
|  | Devprayag | **S4** | 30.1500 | 78.6000 | 830 |
|  | Haridwar | **S5** | 29.9667 | 78.1667 | 314 |
|  | Bijnor | **S6** | 29.3700 | 78.1300 | 225 |
|  | Narora | **S7** | 28.1967 | 78.3814 | 174 |
| Middle stretch | Budaun | **S8** | 28.0500 | 79.1200 | 164 |
|  | Farrukhabad | **S9** | 27.3700 | 79.6300 | 151 |
|  | Bithoor | **S10** | 26.6127 | 80.2719 | 126 |
|  | Dalmau | **S11** | 26.0700 | 81.0300 | 115 |
|  | Prayagraj | **S12** | 25.4500 | 81.8500 | 98 |
|  | Mirzapur | **S13** | 25.1500 | 82.5800 | 80 |
|  | Varanasi | **S14** | 25.3189 | 83.0128 | 81 |
|  | Ballia | **S15** | 25.7625 | 84.1519 | 67 |
| Lower stretch | Revelganj | **S16** | 25.7800 | 84.6700 | 52 |
|  | Patna | **S17** | 25.6100 | 85.1414 | 53 |
|  | Barh | **S18** | 25.4800 | 85.7200 | 47 |
|  | Bahachouki | **S19** | 25.2975 | 86.3561 | 55 |
|  | Farka | **S20** | 25.2287 | 87.0949 | 42 |
|  | Sahebganj | **S21** | 25.2500 | 87.6500 | 16 |
|  | Farakka<br>Barrage<br>Township | **S22** | 24.8200 | 87.9000 | 30 |
|  | Murshidabad | **S23** | 24.1800 | 88.2700 | 18 |
|  | Mayapur | **S24** | 23.4250 | 88.3908 | 11 |
|  | Hooghly | **S25** | 22.9089 | 88.3967 | 9 |
|  | Gangasagar | **S26** | 22.1910 | 88.1905 | 4 |

### 2.2. Analytical Methods

The methods involve in-situ measurements directly at the water source and laboratory-based analyses for more comprehensive assessments. Key physicochemical parameters, such as dissolved oxygen, biochemical oxygen demand, turbidity, total hardness, and total coliform, are determined through established protocols shown in Table.

Various parameters, including temperature, pH, turbidity, electrical conductivity, dissolved oxygen, total solids, Nitrate-N, sodium, potassium, biochemical oxygen demand (BOD), chemical oxygen demand (COD), alkalinity, total hardness, chloride, sulfate, fluoride, and total coliform, were analyzed using specific methods as detailed in the CPCB, World Health Organization (WHO), and Bureau of Indian Standards (BIS) guidelines. These methods encompassed color comparison, odor assessment, thermometer readings, pH meter measurements, nephelometer/turbidimeter readings, conductivity meter assessments, DO meter or Winkler modified method for dissolved oxygen, gravimetric analysis for total solids, spectrophotometry for nitrate-N, flame photometry for sodium and potassium, BOD measurement, potassium dichromate method for COD, titrimetric analyses for alkalinity, total hardness, and carbonate as CaCO3, argentometric titration for chloride, turbidimetry for sulphate, ion meter and calorimetry for fluoride, and the Most Probable Number (MPN) Method for total coliform enumeration, providing a comprehensive overview of water quality across multiple parameters.

### 2.3. Suitability Analysis

Suitability analysis, a technique within Geographic Information Systems (GIS), aims to assess the suitability of diverse locations for particular activities, developments, or land uses. By evaluating a range of spatial data layers encompassing factors like land cover, terrain, infrastructure, environmental limitations, socio-economic variables, and regulatory constraints, this method identifies the most fitting or advantageous locations for specific purposes. For assessing the suitability of drinking water quality (NSFWQI), evaluating the suitability for irrigation (SAR), and determining the suitability for industrial purposes (LSI), various analyses are conducted. These analyses encompass a range of factors specific to each purpose, aiding in the assessment of suitability.

#### 2.3.1. Drinking Water

The National Sanitation Foundation Water Quality Index was used in this study to determine the acceptability of drinking water from the Ganga River. The NSF-WQI is a generally established instrument for assessing the safety and quality of drinking water, particularly along the Ganga. This index takes into account a variety of elements known to influence water quality, such as temperature, pH, turbidity, total dissolved solids (TDS), and concentrations of pollutants such as heavy metals and pathogens.

### The NSF-WQI was calculated using the formula: NSF-WQI = Wi*Pi/Wi

Where,

Wi=weighting coefficient for the ith parameter Pi=is the value of the ith parameter

The estimated NSF-WQI values were divided into ranges to represent water quality. These categories were based on the standards specified by Brown et al. (1970), who classified water quality as entirely totally unsuitable (0-25), unsuitable (25-50), lower medium suitable (50-60), medium suitable (60-75), suitable (70-90), and totally suitable (90-100).

#### 2.3.2. Irrigation Water

The Ganga River’s water suitability for irrigation was evaluated using the Sodium Adsorption Ratio (SAR), an important statistic for analyzing potential dangers connected with salts in irrigation water. SAR calculates the ratio of sodium to calcium and magnesium ions, which serves as an indicator of sodium risks in water. This ratio provides useful information about the extent of cation exchange interactions that irrigation water may experience within the soil. High salt levels have the potential to penetrate soil particles, altering soil properties and reducing permeability (Interpreting Water Quality Test Results, First Edition NSW DPI Agriculture Water Unit, 2016; Ayers and Bronson, 1975).

The SAR was calculated using the formula:

### SAR = Na+/ (Ca2+ + Mg2+ /2)1/2

Water samples were analyzed to determine concentrations of sodium, calcium, and magnesium ions, essential for SAR calculation. SAR values were categorized into suitability classifications based on thresholds established by Haritash et al. (2016), defining suitability as: Suitable (<10), Medium suitable (10-18), Unsuitable (18-26), and unsuitable (>26).

#### 2.3.3. Industrial water

Industrial water, utilized for various purposes such as cooling systems, boiler feedwater, and manufacturing processes, requires meticulous monitoring to prevent pipe corrosion within utilities. Saturation indices, notably the Langelier Saturation Index (LSI), play a crucial role in evaluating water’s potential to either precipitate or dissolve calcium carbonate, essential for maintaining water quality standards in industrial settings.

The LSI is a calculated value used to predict the stability of calcium carbonate in water. It is determined by the difference between the measured system pH and the saturation pH:

### LSI = pH (measured)-pHs

The LSI provides insights into whether water will precipitate, dissolve, or maintain equilibrium with calcium carbonate. A negative LSI indicates water’s potential to dissolve calcium carbonate, while a positive LSI suggests the likelihood of scale formation. An LSI close to zero implies borderline scale potential, emphasizing the critical point of water quality where external factors such as temperature, evaporation, and water chemistry may influence the outcome. According to Haritash et al. (2016), LSI classifications include Suitable (negative), Medium suitable (0), and Unsuitable (positive) categories.

### 2.5. Statistical Analysis

The statistical analyses were performed using GraphPad Prism and SPSS 25. Descriptive statistics, including the maximum, minimum, median, and standard deviation, were calculated for each set. The normality of the probability distributions of heavy metals was assessed using the Shapiro-Wilk test. The differences between the median values among the UGS, MGS, and LGS were evaluated using the Kruskal-Wallis test at a 95% confidence interval. The p-value of less than 0.05 was considered significant. To identify the sources of pollution, multivariate analyses such as principal component analysis (PCA) and cluster analysis were conducted. The collected data for PCA were first standardized as Z values to ensure a uniform range. PCA was used to assess the continuous presence of individual metals in the samples, and the highest contribution of each metal was examined. Cluster analysis using Pearson correlation was also carried out to validate the PCA results. The application of PCA in the assessment of heavy metal pollution is a well-established method, allowing for the identification of sources and the overall assessment of the interaction of variables [3].

## 3. Results and Discussion

### Physico-biochemical characteristics of water of River Ganga in India

The physical-biochemical characteristics of the water in the River Ganga, are of immense significance due to their impact on the river’s overall health and the well-being of the communities dependent on it. The river exhibits a complex interplay of physical and biochemical factors that contribute to its overall water quality. Physically, the Ganga is known for its diverse flow patterns, ranging from turbulent stretches to calmer regions, influencing sediment transport and distribution. The river’s temperature varies across its course, affecting the dissolved oxygen levels crucial for aquatic life. Biochemically, the River Ganga sustains a rich biodiversity and supports a variety of aquatic ecosystems. The water carries essential nutrients and minerals vital for the growth of aquatic flora and fauna. However, the river faces challenges such as high levels of organic and inorganic pollutants, including industrial effluents and agricultural runoff, which can significantly impact its biochemical composition.

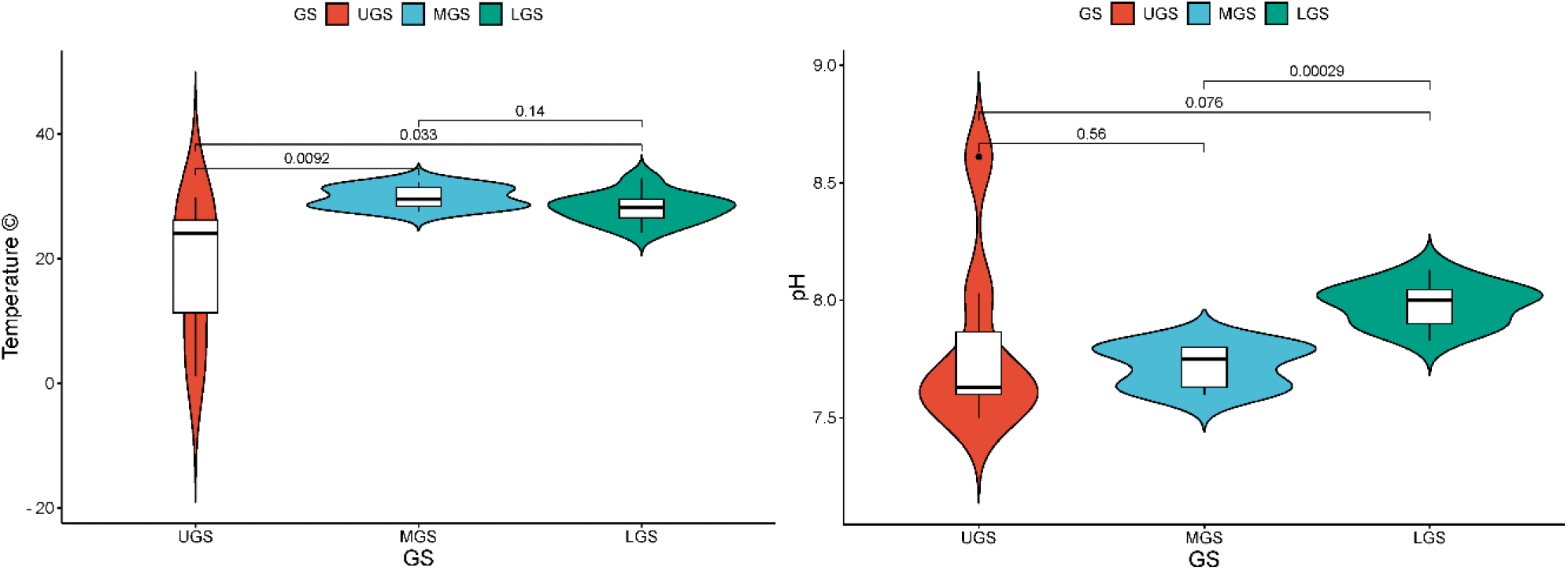
Plot of calculated values of temperature and pH of Ganga river water samples across upper, middle, and lower stretch of Ganga river

The temperature in the upper stretch of the river varied from a freezing 1.1°C to 29.8°C at Narora (S7), with turbidity levels ranging from 2.02 to 37.53 NTU, beginning at Gomukh (S1) and Gangotri (S2), the river’s source. The pH ranged from 7.5 to 8.16, remaining largely constant. The middle stretch saw a minor increase in temperature, with measurements ranging from 27.6°C to 32.2°C. The range of turbidity levels was larger, ranging from 4.66 to 61.31 NTU, suggesting increasing sedimentation and human activities. The conditions were somewhat alkaline, as indicated by pH values ranging from 7.6 to 7.86. Ultimately, temperatures in the lower stretch were similar to those in the middle stretch, but turbidity levels varied greatly, peaking at Diamond Harbour at 694.3 NTU. In general, the pH ranged from 7.83 to 8.16, which is higher.

**Fig. 2.**
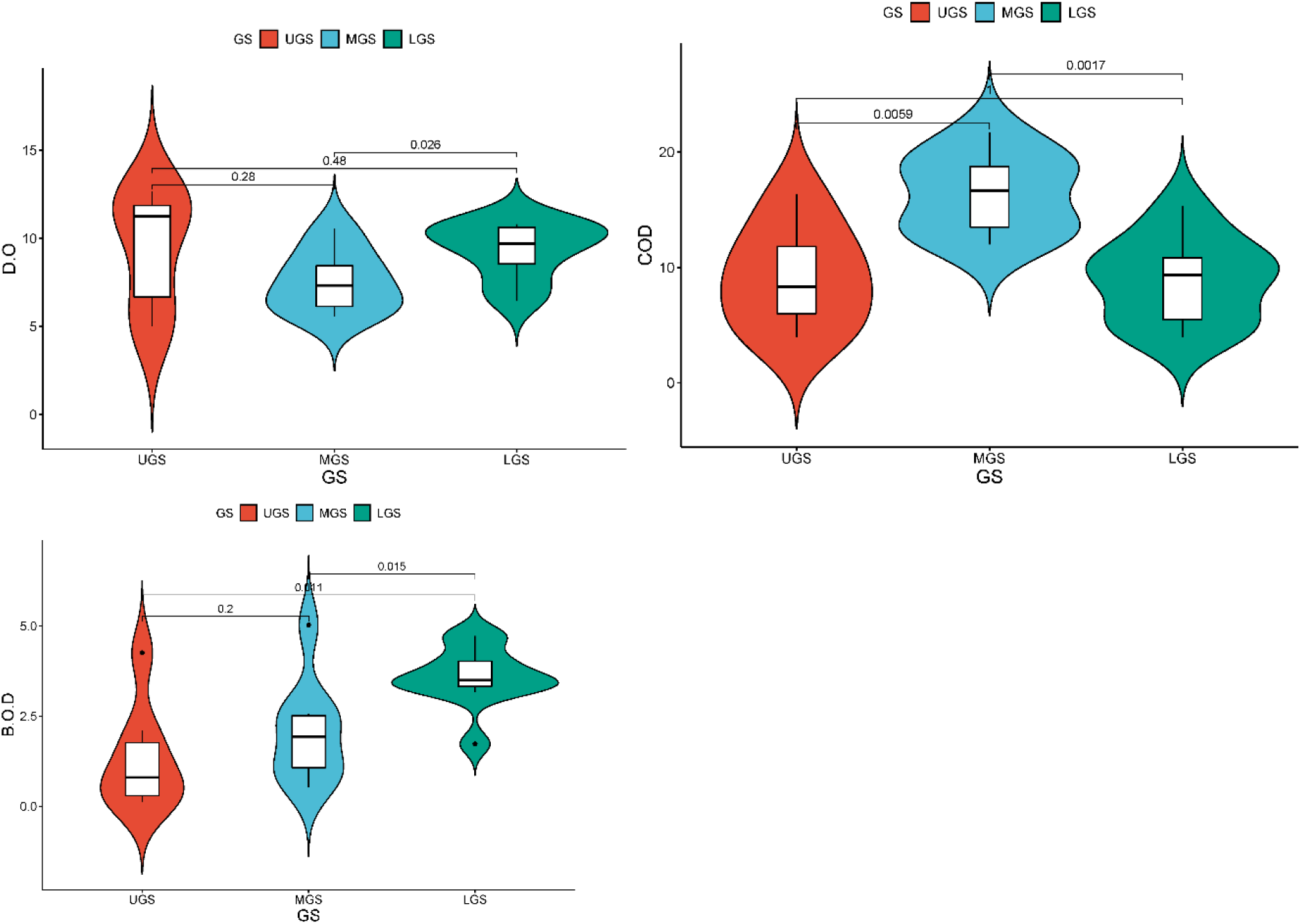
Plot of calculated values of DO, COD and BOD of Ganga river water samples across upper, middle and lower stretch of Ganga river

Dissolved oxygen (DO) values in the upper Ganga stretch were 8.2 mg/L at Devprayag (S4) and 12.66 mg/L at Gomukh (S1). Haridwar (S5) had the greatest levels of chemical oxygen demand (COD) at 16.33 mg/L, while Gomukh (S1) had the lowest levels at 4 mg/L. Meanwhile, biochemical oxygen demand (BOD) peaked at 4.26 mg/L and was lowest in Gangotri (S2) at 0.13 mg/L. As one moved to the middle Ganga section, significant differences continued. At 9.3 mg/L, Budaun (S8) had the highest DO levels, while Farrukhabad (S9) had the lowest, at 5.56 mg/L. With 1.36 mg/L, Dalmau (S11) had the highest BOD levels, whereas Budaun (S8) had the lowest, at 2.5 mg/L. With 19 mg/L, Prayagraj (S12) had the highest COD readings, whereas Budaun (S8) had the lowest, at 12 mg/L. Finally, a variety of tendencies started to appear in the lower Ganga stretch. Diamond Harbour (S26) had the lowest DO levels at 6.96 mg/L, while Revelganj (S16) had the highest at 10.8 mg/L. Revelganj (S16) had the highest BOD values at 4.73 mg/L, while Patna (S17) had the lowest at 3.6 mg/L. Barh (S18) had the lowest COD levels at 4.67 mg/L, while Sahebganj (S21) had the highest at 15.33 mg/L.

**Fig. 2.**
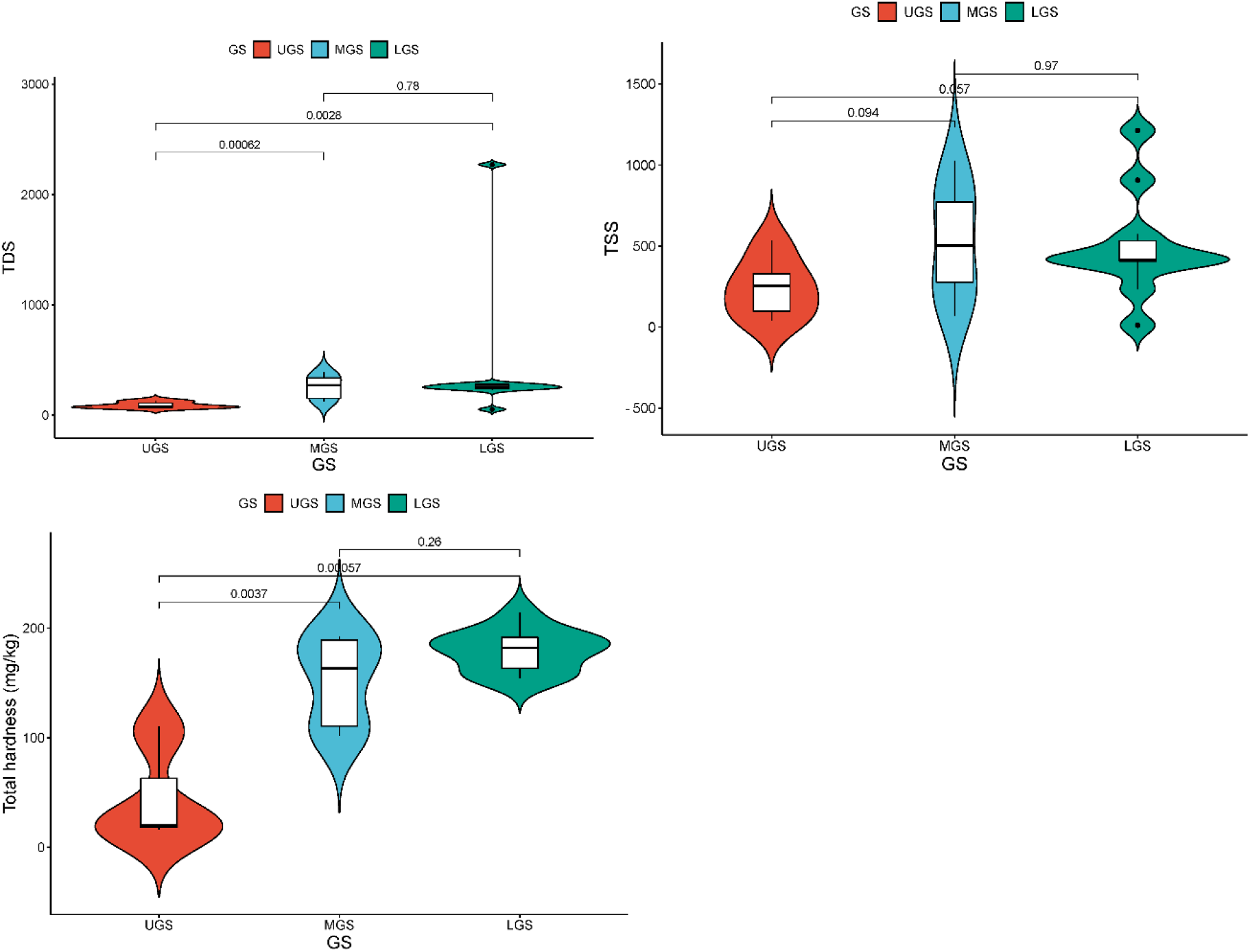
Plot of calculated values of TDS, TSSS and total hardness of Ganga river water samples across upper, middle and lower stretch of Ganga river

With a Total Dissolved Solids (TDS) concentration of 134.93 mg/kg, site Narora (S7) in the upper Ganga River segment showed the highest amount of dissolved substances in the water. On the other hand, site Gangotri (S2) had the lowest TDS content in this length, at 66.54 mg/kg. Site Bijnor (S6) had the highest Total Suspended Solids (TSS) value of 533.33 mg/kg, suggesting that there were a significant number of suspended particles in the water. Site Gangotri (S2) had the lowest TSS concentration, measuring 39.33 mg/kg. Site Uttarkashi (S3) had the lowest Total Hardness at 16 mg/kg, while Site Bijnor (S6) had the highest value at 110 mg/kg, indicating rather hard water. As we move down the middle stretch of the river, site Bithoor (S10) has the greatest TDS concentration, 157.13 mg/kg, while site Farrukhabad (S9) has the lowest—127.86 mg/kg. Similar to this, site Bithoor (S10) had the greatest TSS concentration at 1026.67 mg/kg, whereas site Mirzapur (S13) had the lowest TSS concentration at 66.67 mg/kg. When it came to total hardness, site Varanasi (S14) had the greatest value—192 mg/kg—while site Farrukhabad (S9) had the lowest—102 mg/kg. With a remarkably high value of 2271 mg/kg, site Gangasagar (S26) in the lower Ganga River segment displayed the highest TDS concentration, suggesting a substantial presence of dissolved chemicals. In the lower stretch, site Hooghly (S25) had the lowest TDS content in this length, at 53.06 mg/kg. Site Patna (S17) had the highest Total Suspended Solids value of 1213.00 mg/kg, indicating a significant number of suspended particles in the water; site Hooghly (S25) had the lowest TSS concentration of 11.55 mg/kg. In terms of total hardness, site Bahachouki (S19) had the greatest value (214 mg/kg), whereas site Mayapur (S24) had the lowest value (154 mg/kg).

**Fig. 2.**
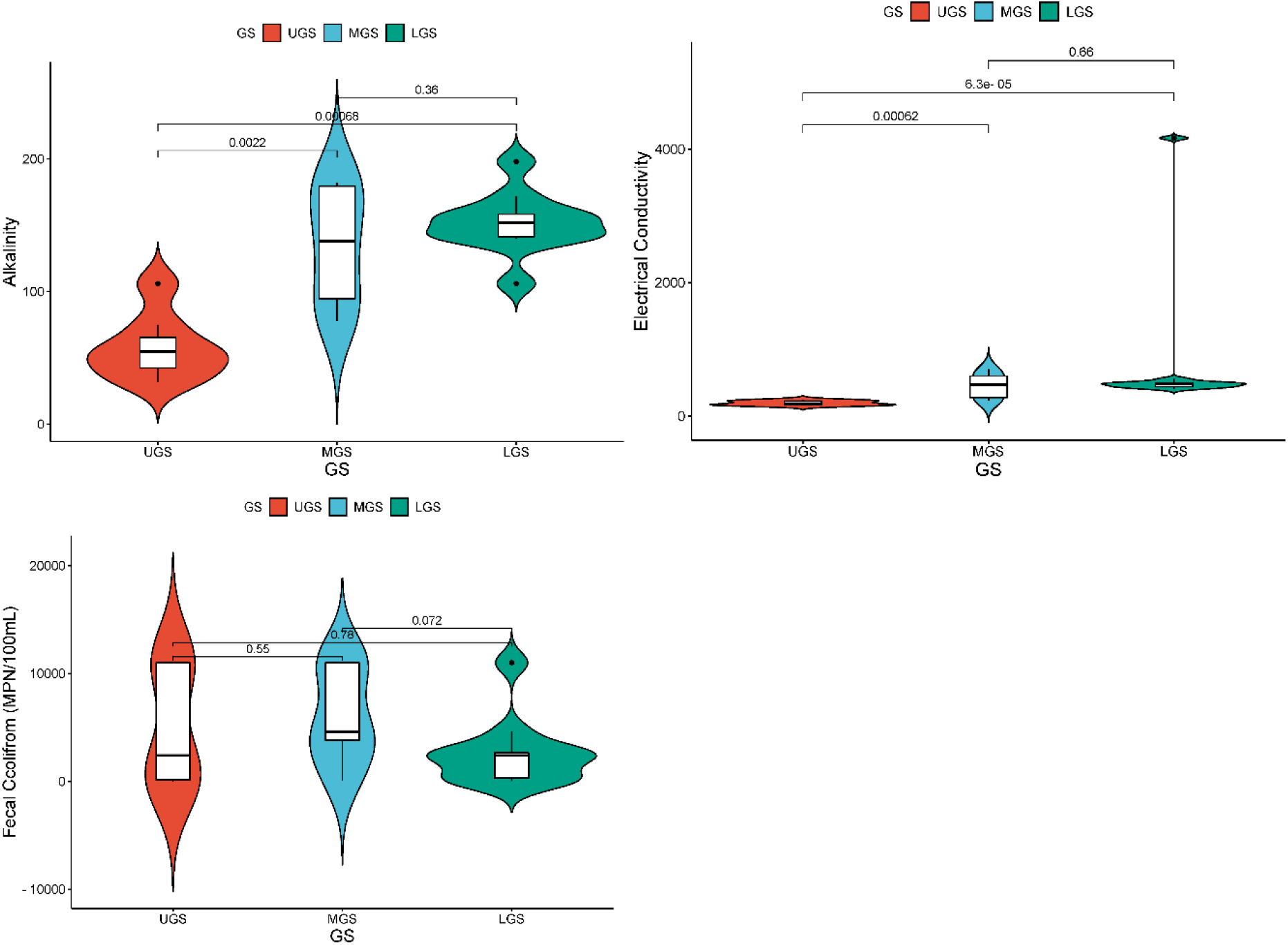
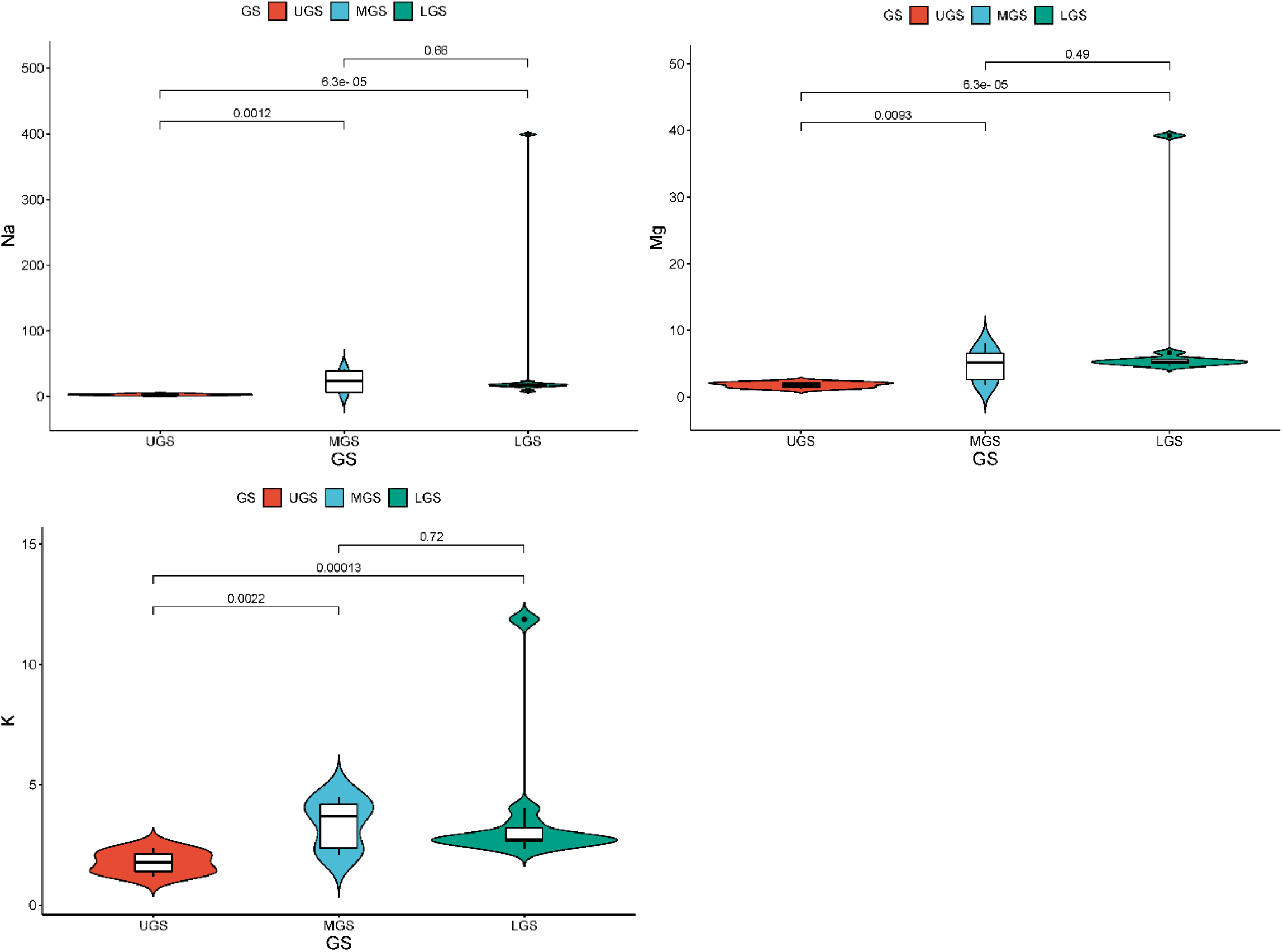
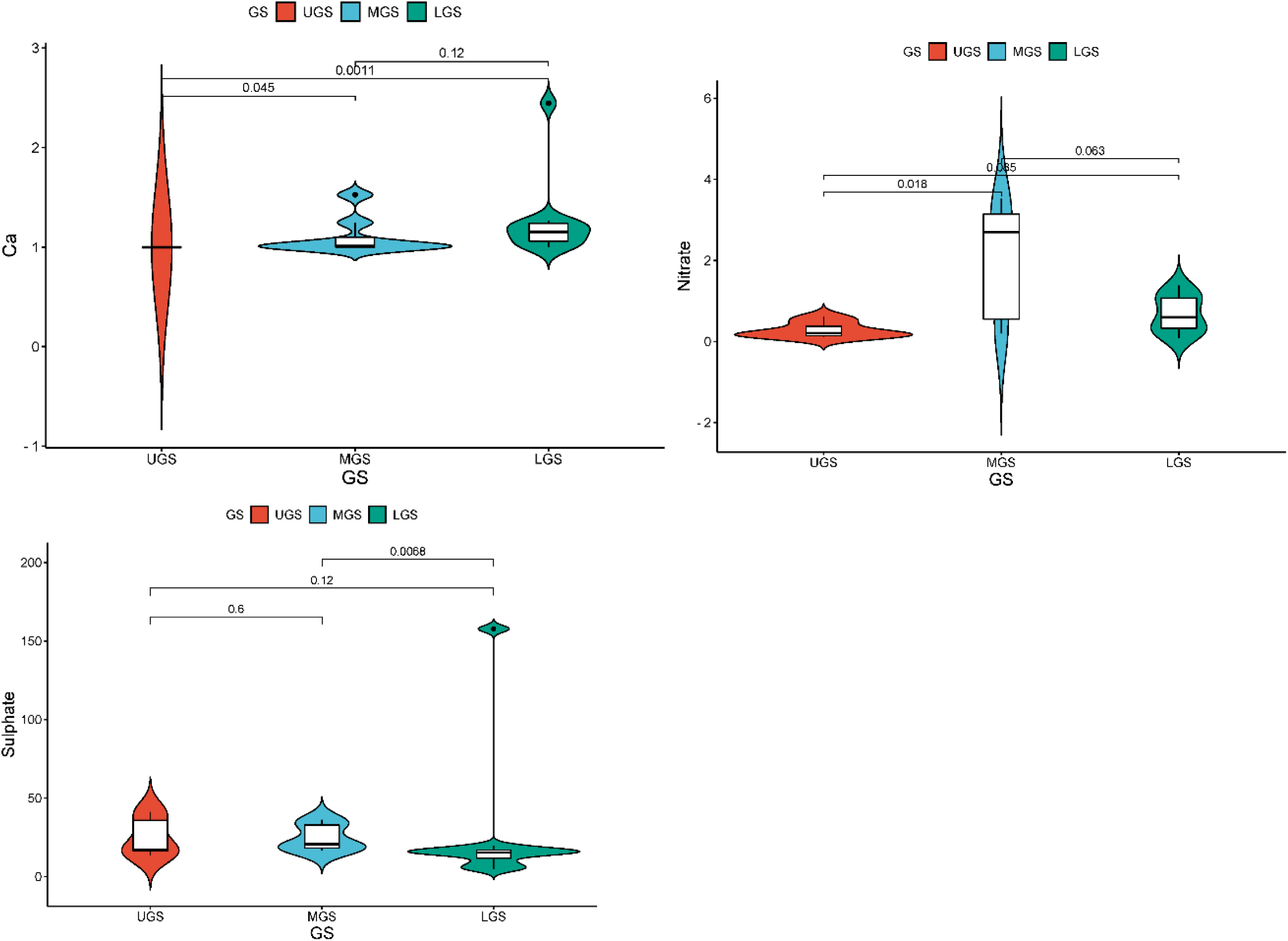
Plot of calculated values of alkalinity, electrical conductivity, and Fecal coliform of Ganga river water samples across upper, middle, and lower stretch of Ganga river

**Fig. 2.**
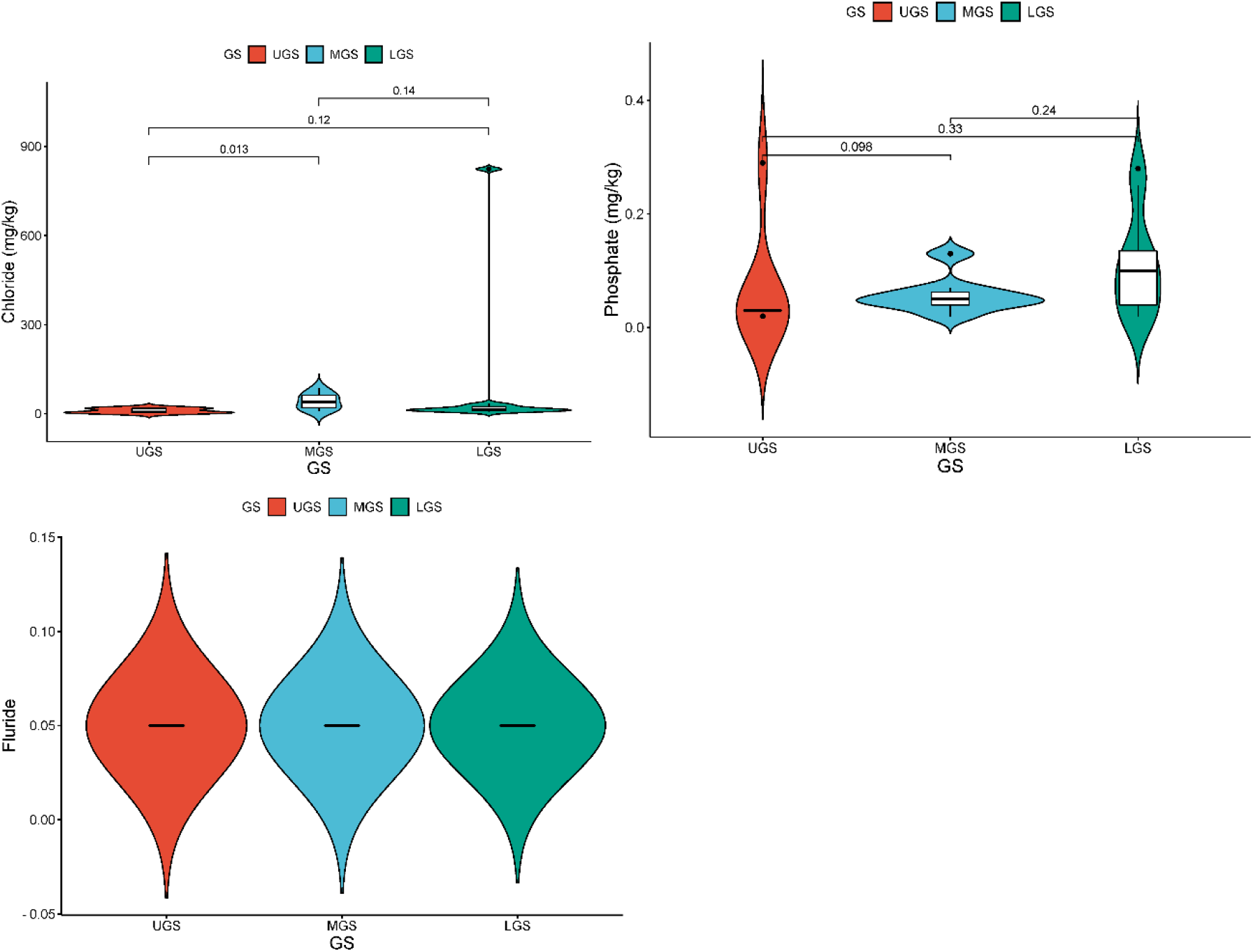
Plot of calculated values of sodium, magnesium, potassium, calcium, nitrate, sulfate, chloride, phosphate, and fluoride of Ganga river water samples across upper, middle and lower stretch of Ganga river

In the upper stretch, the recorded alkalinity levels range from a minimum of 32.00 mg/L at Gaumukh (S1) to a maximum of 106.00 mg/L at Narora (S7). Similarly, the electrical conductivity in this segment varies from a minimum of 168.80 μS/cm at Gaumukh (S1) to a maximum of 244.26 μS/cm at Narora (S7). Moving to the middle stretch, the alkalinity levels show a range between 78.00 mg/L at Budaun (S8) and 180.00 mg/L at Ballia (S15). Conversely, the electrical conductivity exhibits a wider range, with values spanning from 248.03 μS/cm at Budaun (S8) to 701.16 μS/cm at Ballia (S15). Finally, in the lower stretch, the alkalinity ranges from 106.00 mg/L at Gangasagar (S26) to 198.00 mg/L at Bahachouki (S19). Notably, the electrical conductivity in this stretch displays significant variation, with a minimum value of 408.46 μS/cm at Revelganj (S16) and a maximum value of 4167.00 μS/cm at Gangasagar (S26). The upper, middle, and lower Ganga Rivers exhibit notable differences in fecal coliform (FC) levels. FC concentrations in the upper stretch vary from 0 to 11000 MPN/100 mL, with Gaumukh (S1), Devprayag (S4), and Bijnor (S6) recording the highest value of 11000 MPN/100 mL and Gaumukh (S1) displaying the lowest value of 0 MPN/100 mL FC. With Budaun (S8) recording the lowest value of 94 MPN/100 mL and Bithoor (S10), Mirzapur (S13), and Varanasi (S14) documenting the highest value of 11000 MPN/100 mL, the middle stretch FC levels vary between 94 and 11000 MPN/100 mL. With Revelganj (S16) registering the lowest value of 30 MPN/100 mL and Gangasagar (S26) recording the highest value of 11000 MPN/100 mL, the lower stretch FC concentrations vary from 30 to 11000 MPN/100 mL.

Within the upper stretch, the following average concentrations were found: 2.867 mg/kg, 1.791 mg/kg, 1.783 mg/kg, 1.000 mg/kg, 0.277 mg/kg, 22.188 mg/kg, 13.231 mg/kg, 0.026 mg/kg, and 0.050 mg/kg for sodium, magnesium, potassium, calcium, nitrate, sulfate, chloride, phosphate, and fluoride, respective. The concentrations of sodium, magnesium, potassium, calcium, nitrate, sulfate, chloride, and phosphate increase significantly as one moves to the middle stretch with average values of 25.948 mg/kg, 5.474 mg/kg, 3.199 mg/kg, 1.200 mg/kg, 1.955 mg/kg, 27.264 mg/kg, 46.586 mg/kg, and 0.081 mg/kg, respectively. Lastly, the concentrations are highest in the lower stretch where the mean values of calcium, nitrate, sulfate, chloride, phosphate, sodium, magnesium, potassium, and fluoride are 45.678 mg/kg, 8.598 mg/kg, 3.142 mg/kg, 1.170 mg/kg, 1.081 mg/kg, 98.264 mg/kg, 164.092 mg/kg, 0.105 mg/kg, and 0.050 mg/kg, respectively.

### 3.2. Suitability of Drinking Water

To check the suitability of Ganga river water for drinking, several factors were taken into consideration. These comprised the following: fecal coliform (FC) levels, phosphate, nitrate, temperature, pH, turbidity, dissolved oxygen (DO), and biochemical oxygen demand (BOD). The detailed data across upper, middle and lower stretch of Ganga river of each parameters is shown below Figure

**Fig. 2:**
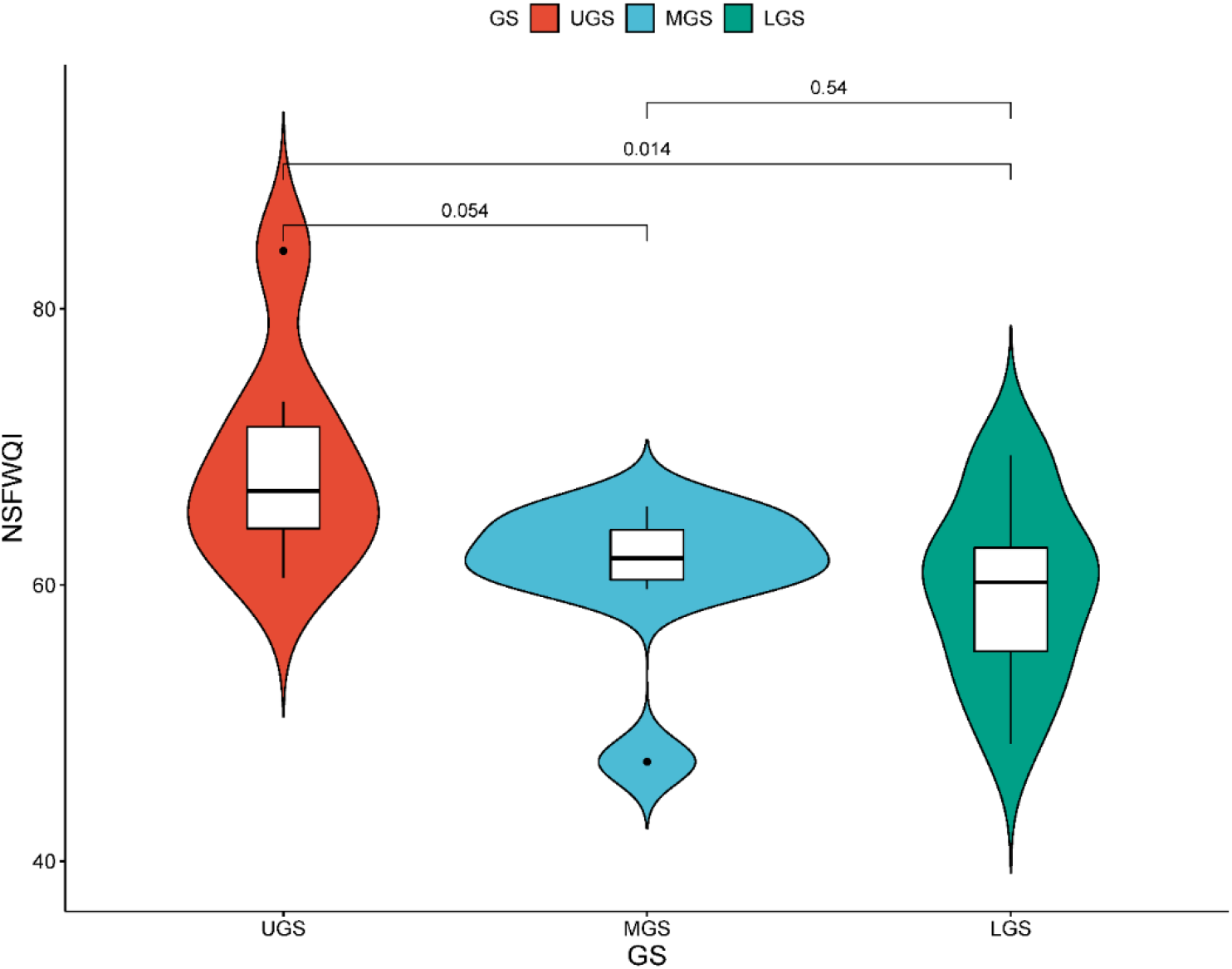
Plot of National Sanitation Foundation water quality index across. upper, middle and lower stretch of Ganga river

The data presented indicates variations in water quality along different stretches of the Ganges River. The Upper Ganges Stretch (UGS) generally exhibits higher quality water, with values ranging from 84.2 to 60.5, suggesting relatively stable conditions. In contrast, the Middle Ganges Stretch (MGS) displays slightly lower water quality, ranging from 65.7 to 47.2, indicating some variability and potential concerns. The Lower Ganges Stretch (LGS) shows the widest range of values, from 68.6 to 48.5, with fluctuations indicating less consistent water quality, potentially raising more significant environmental concerns. Overall, while the UGS demonstrates the best water quality, the MGS and LGS exhibit lower quality water with increasing variability and occasional drops, highlighting potential challenges in maintaining water quality standards along the Ganges River.

### 3.3. Suitability of Irrigation water

The suitability of Ganga River water for irrigation, reveals promising attributes based on key parameters such as sodium (Na), calcium (Ca), and magnesium (Mg). The detailed data across **the** upper, middle, and lower stretch of the Ganga river of each parameter is shown below figure.

**Fig. 3:**
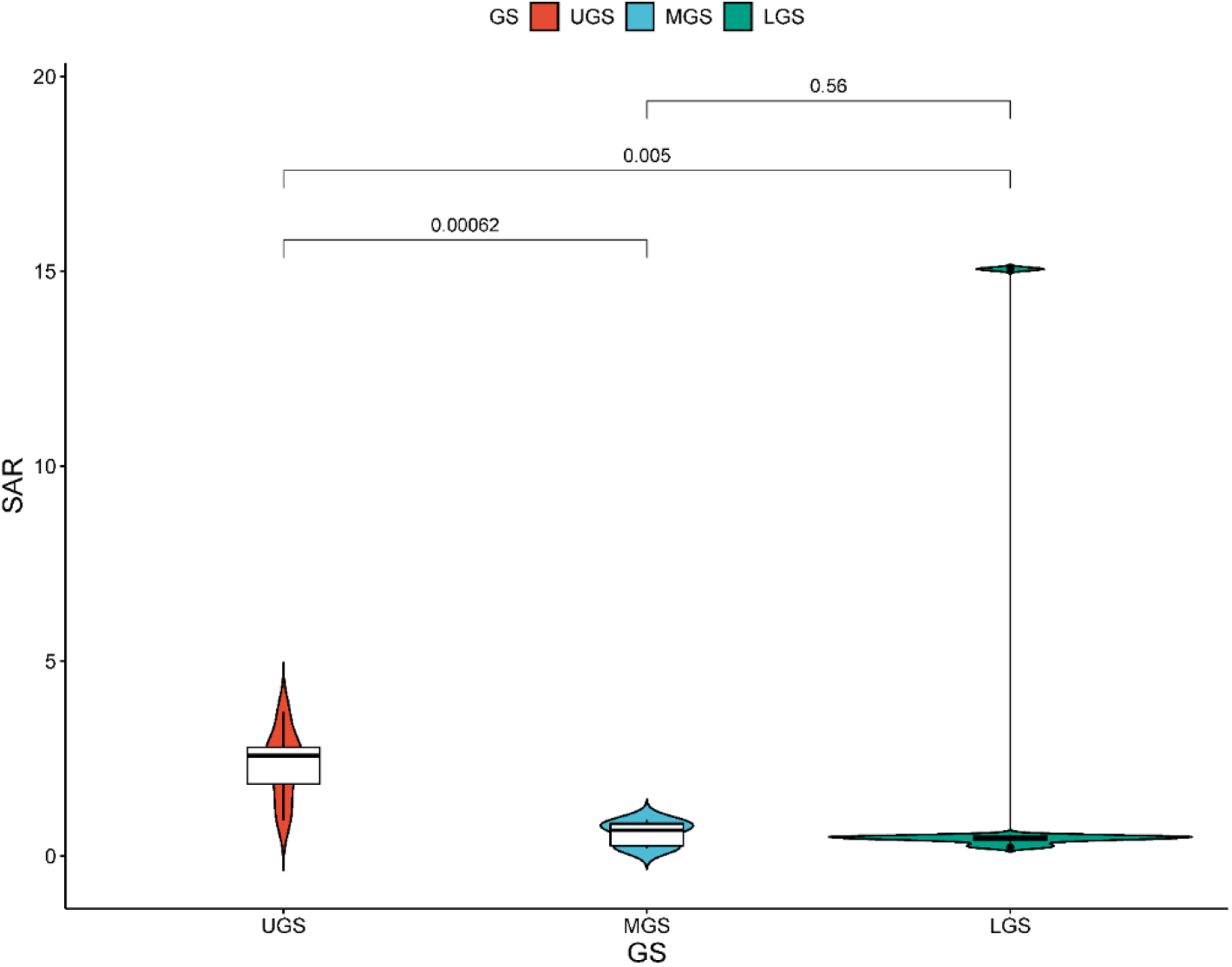
Plot of Sodium Absorption Ratio index across. upper, middle and lower stretch of Ganga river

In this study, SAR was found in the range of 0.91–87.43. The data illustrates the Suspended Ammonia Ratio (SAR) across three segments of the Ganges River the Upper Ganges Stretch (UGS), Middle Ganges Stretch (MGS), and Lower Ganges Stretch (LGS). In the UGS, SAR values range from 0.91 to 3.69, indicating moderate variability, while the MGS shows higher levels, with SAR values ranging from 3.54 to 20.04, and the LGS displays the highest variability and levels, ranging from 4.36 to an alarming 87.43. Overall The values of parameters utilized in SAR and the calculated values provided in the table have been provided in Table 3. These findings suggest that while all stretches exhibit some level of suspended ammonia, the LGS faces the most significant challenges, warranting immediate environmental intervention to mitigate pollution levels along this section of the Ganges River.

### 3.3. Suitability of Industrial water

The suitability of industrial water in the Ganga River was assessed based on several key parameters, including alkalinity, electrical conductivity, calcium hardness, temperature, and pH. The detailed data across the upper, middle, and lower stretch of the Ganga river of each parameter is shown below figure

**Fig. 5:**
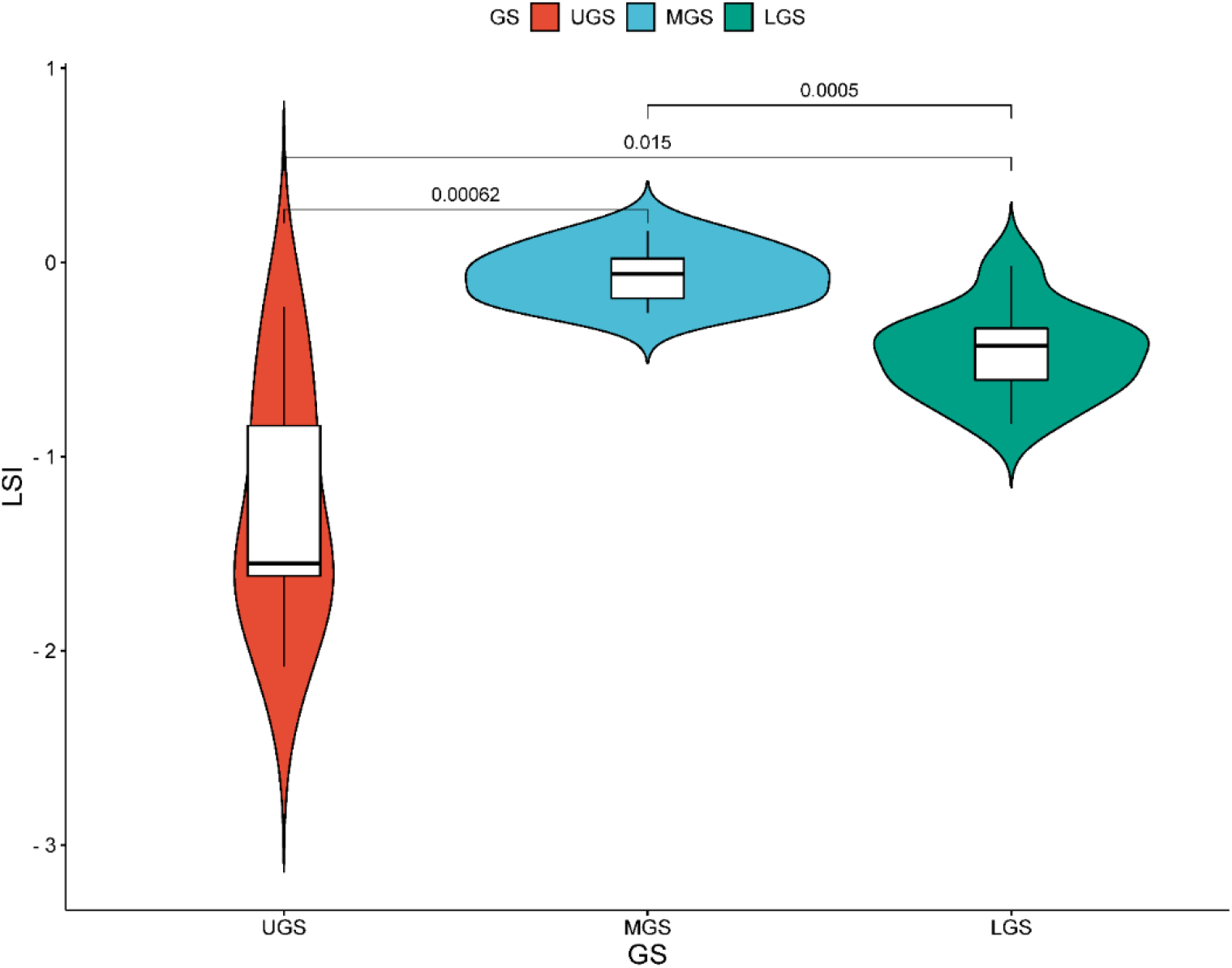
Plot of Langelier Saturation Index across. upper, middle and lower stretch of Ganga river

The Langelier saturation index (LSI) values across the Upper Ganges Stretch (UGS), Middle Ganges Stretch (MGS), and Lower Ganges Stretch (LGS) indicate varying levels of pollution. In the UGS, LSI ranges from -2.08 to -0.23, with more negative values suggesting higher pollution levels. Conversely, the MGS displays a narrower range of -0.26 to 0.16, indicating relatively cleaner water conditions compared to the UGS. The LGS exhibits a similar range of variability to the UGS, with LSI values ranging from -0.83 to -0.02. Overall, the UGS tends to have higher pollution levels, while the MGS shows slightly cleaner water conditions, and the LGS demonstrates variability from relatively high to lower pollution levels. Overall The values of parameters utilized in LSI and the calculated values provided in the table have been provided in Table 4. These findings underscore potential environmental concerns, particularly in the Upper Ganges Stretch, necessitating remediation efforts to improve water quality along this segment of the Ganges River.

### 3.4. Classification of water quality for drinking, irrigation, and industrial

| Quality | NSFWQI | SAR | LSI |
| --- | --- | --- | --- |
| Totally unsuitable | 0-25 | >26 | - |
| unsuitable | 25-50 | 18-26 | +ve value |
| lower medium suitable | 50-60 | - | - |
| medium suitable | 6-75 | 10-18 | 0 |
| suitable | 70-90 | <10 | -ve value |
| Totally suitable | 90-100 | - | - |

## 3. Conclusion

In evaluating the suitability analysis of Gangetic water from the Upper, Middle, and Lower Ganga Rivers, utilizing parameters such as NSFWQI, SAR, and LSI, several key observations were made.

Firstly, the study revealed that the water quality of the Ganga River was generally good in the Upper segment, particularly in locations around Rishikesh town. Here, the water exhibited qualities conducive to drinking purposes with minimal treatment post-disinfection. Additionally, heavy metal concentrations were either non-detectable or within safe limits, further affirming the suitability of the water for consumption. However, as the study progressed downstream towards the Middle and Lower segments of the Ganga River, higher levels of pollutants were observed, particularly in Total Dissolved Solids (TDS), organic matter, and Most Probable Number (MPN). Despite this, these areas may still serve as suitable locations for organized outdoor bathing activities for tourists and pilgrims, albeit with necessary precautions. The notable concern was issues related to night soil disposal in the riverbed and wastewater discharge through open channels, which necessitated immediate attention to control MPN and organic load. Regular monitoring and stringent measures were recommended to address MPN, a critical parameter requiring ongoing vigilance.

Furthermore, the addition of phosphate from wastewater channels posed a risk of eutrophication, especially during lean flow periods. To mitigate this risk, channels could be intercepted and diverted to sewage treatment plants to prevent phosphate contamination. Despite these challenges, the overall water quality was observed to be good to excellent, particularly in terms of suitability for irrigation over prolonged periods. However, in industrial applications, while the water would not lead to scale formation, it may result in pipe corrosion. Nevertheless, with minimal pretreatment, the water could still be utilized in various industrial processes. In conclusion, the evaluation highlighted the dynamic nature of water quality along the Upper, Middle, and Lower stretches of the Ganga River. While challenges exist, proactive measures such as regular monitoring, effective waste management practices, and targeted interventions can contribute to the preservation and enhancement of water quality in the Ganga River basin.

